# Longitudinal Investigation of Aortic Dissection in Mice with Computational Fluid Dynamics

**DOI:** 10.1101/2023.04.24.538163

**Authors:** Kathrin Bäumler, Evan H. Phillips, Noelia Grande Gutiérrez, Dominik Fleischmann, Alison L. Marsden, Craig J. Goergen

## Abstract

Patients with aortic dissection require lifelong surveillance to monitor disease progression and detect late adverse events such as aneurysmal dilation, malperfusion or refractory pain. The variety and complexity of aortic dissection have so far eluded definitive predictions of occurrence and timing of late adverse events. The search for early indicators of late adverse events has been based mostly on morphologic features, and one commonly observed risk factor is partial thrombosis of the false lumen. While the effect of partial thrombosis on disease progression is incompletely understood, hemodynamic factors, including low velocity or stagnant flow, are likely to play a role. In this study we investigated the progression of false lumen intramural thrombus formation in four mice with angiotensin II-induced aortic dissection. Based on 3D B-mode ultrasound images, we created segmentations of the diseased aorta including the true lumen, false lumen, and thrombus. These geometries were then used to run computational fluid dynamic simulations with subject-specific boundary conditions. Each mouse was followed for seven days and 4-5 longitudinal image datasets were acquired for each animal. We found that false lumina with a single entry tear tend to have smaller mean relative velocities, and at the same time are subject to a larger false lumen thrombus ratio. Likewise, regions of low velocity correlated with regions of elevated endothelial cell activation potential and higher particle residence times. These findings support the hypothesis that flow stagnation is the predominant hemodynamic factor that results in a large thrombus ratio in false lumina, particularly those with a single entry tear. Additional work will be needed to further explore the intricacies of these complex experimental vascular lesions and how the hemodynamic conditions compare to human aortic dissections.

## Introduction

Aortic dissection is an acute life-threatening abnormality of the aorta in which the medial layer of the aorta is delaminated. As a result, a double channel aorta develops with a true lumen (TL) and a false lumen (FL), separated by a dissection membrane or flap. In most cases, an entrance tear in the dissection flap can be identified by medical imaging such as computed tomography (CT), magnetic resonance imaging (MRI), or ultrasound (US). Re-entrance tears and ad-ditional fenestrations in the dissection flap are often present and enable a communication between the TL and the FL. Thrombosis in aortic dissection is especially critical due to its effects on both short and long-term outcomes. (1) found that partial thrombosis of the FL was an independent predictor of post-discharge mortality in patients with Type B aortic dissection. The same study reported the lowest rate of 3year mortality in patients with a patent FL (13.7 *±* 7.1%), compared to those with a complete thrombosed FL (22.6 *±\* 22.6%) and a partial thrombosed FL (31.6 *±* 12.4%). In a study by (2), the increase in partial FL thrombosis area over time was strongly associated with late adverse events in medically managed patients with Type B aortic dissections (median follow-up time of 4 years with a range of 10 days to 12.7 years). Likewise, (3) and (4) found that a partially thrombosed FL yielded higher aortic growth rates compared to patent or fully thrombosed FL.

In addition to human studies, animal models (5–8) have been used extensively to study the pathophysiology of aortic dissections and advance our understanding of thrombus formation in complex lesions. The angiotensinII (AngII) infusion model (5) has distinct features, including dissection and typically leftward expansion just above the level of the kidneys (5, 7), and separation of the media and adventitia creating a FL. Indeed, as with human aortic dissections, medial tears and thrombosis can be identified (6, 8, 9). In a study by (10), four mice were infused with AngII and developed dissecting suprarenal abdominal aortic aneurysms. The authors introduced a combination of moderate resolution (40 *μ*m) *in vivo* anatomical and Doppler US imaging with high resolution (7 *μ*m) *in vitro* optical coherence tomography (OCT). Hemodynamic simulations in the four mice, of which three developed thrombus in the FL, were then performed. The findings suggested a multi-modality imaging approach could be used to create optimized geometric models for hemodynamic simulations. Simulation results indicated low time-averaged wall shear stress (TAWSS) and presence of vortical flow in the FL when compared to the TL. However only one timepoint was investigated as *in vitro* OCT imaging was used, negating the possibility of longitudinally following the same animal over time.

Based on these same mouse models, (11) developed a numerical framework to predict thrombus formation and growth in aortic dissections. Their model includes a two-way coupling of activated and resting platelets on the flow field, where platelets are treated as particles that interact with each other and the arterial wall. The actual location of thrombus formation was determined based on a volume fraction of the activated platelets and fibrinogen. The authors found prominent effects of hemodynamic quantities such as shear rates on the final size and shape of thrombus, all while achieving generally good agreement with imaging data.

In the present study, we investigated three murine specimens from (10) and one additional mouse (acquired during the same study with large and complex FL size and morphology) over multiple timepoints up to seven days. We analyzed thrombus growth over time based on US imaging data and performed rigid wall computational fluid dynamic simulations in these mouse models to identify possible correlation between thrombus growth and hemodynamic factors. All mice exhibited rapidly changing thrombus size and location during the observed time period, suggesting a highly complex and dynamic thrombotic process. Our results confirmed that FL with a single opening yielded larger thrombus ratios and lower mean velocities compared to FL with multiple openings. These results support the hypothesis that slow or stagnating flow constitutes the predominant factor in FL thrombosis in the investigated specimen. Hemodynamic results also highlighted increased endothelial cell activation potential, particle residence times, thrombus formation potential, and slow and stagnating flow in FL with a single tear or opening throughout the observation period. In FL with two or more openings, regions of the FL with higher flow ratios exhibited reduced endothelial cell activation potential, particle residence times, and thrombus formation potential.

## Methods

### Mouse model and *in vivo* imaging

In a prior study (10), four male ApoE^−*/*−^ mice were subcutaneously implanted with mini-osmotic pumps loaded with angiotensin II (1000 ng/kg/min) to induce an aortic dissection. Approval to conduct all animal procedures was received from the Purdue University Institutional Animal Care and Use Committee. The suprarenal abdominal aorta (SAA) was monitored every 48 hours post-operatively using US to detect the initiation of a dissection, after which lesion geometry and blood flow were quantified using a variety of US imaging techniques (B-mode, M-mode, pulsed wave (PW) Doppler, and ECG-gated kilohertz visualization (EKV) mode) every two days for seven days (MS550D transducer, Vevo2100, FUJIFILM VisualSonics Inc.).

### *In vivo* pressure data

Diastolic and systolic arterial blood pressures were measured in conscious mice using a tail-cuff device (Coda two-channel, Kent Scientific). Measurements were taken at baseline, on days 3 and 7 post-surgery, and on day 4 post-diagnosis of a lesion. These pressure data informed the numerical simulations by providing target pressures. Since pressure measurement days did not always coincide with simulation days, we applied the latest available post-diagnosis measurement as input for the simulation (Table 1).

**Table 1.** Pressure Tuning. MAP = mean arterial pressure. Target pressure values for numerical simulations are obtained from tail cuff measurements.

|  |  | Targets |  |  | Simulation Results |  |  | Error |  |  |
| --- | --- | --- | --- | --- | --- | --- | --- | --- | --- | --- |
| | | $P_{dia}$ | $P_{sys}$ | MAP | $P_{dia}$ | $P_{sys}$ | MAP | $P_{dia}$ | $P_{sys}$ | MAP |
| M2 | Day 0 | 126.1 | 167.8 | 139.7 | 130.7 | 173.4 | 147.0 | 3.68% | 3.33% | 5.22% |
|  | Day 1 | 126.1 | 167.8 | 139.7 | 132.8 | 174.7 | 149.6 | 5.30% | 4.08% | 7.08% |
|  | Day 3 | 126.1 | 167.8 | 139.7 | 122.7 | 161.9 | 141.5 | 2.72% | 3.56% | 1.27% |
|  | Day 5 | 136.7 | 181.9 | 151.4 | 139.7 | 182.9 | 159.6 | 2.15% | 0.57% | 5.45% |
|  | Day 7 | 136.7 | 181.9 | 151.4 | 130.1 | 173.0 | 148.6 | 4.82% | 4.87% | 1.84% |
| M3 | Day 0 | 161.7 | 199.9 | 174.4 | 168.8 | 209.7 | 187.5 | 4.42% | 4.91% | 7.50% |
|  | Day 1 | - | - | - | - | - | - | - | - | - |
|  | Day 3 | 161.7 | 199.9 | 174.4 | 160.7 | 197.3 | 179.1 | 0.58% | 1.31% | 2.71% |
|  | Day 5 | 161.7 | 199.9 | 174.4 | 161.9 | 203.9 | 181.3 | 0.16% | 2.01% | 3.96% |
|  | Day 7 | 161.7 | 199.9 | 174.4 | 162.6 | 202.1 | 181.7 | 0.59% | 1.12% | 4.16% |
| M4 | Day 0 | - | - | - | - | - | - | - | - | - |
|  | Day 1 | 106.1 | 150.2 | 120.4 | 110.4 | 146.5 | 124.4 | 4.07% | 2.47% | 3.35% |
|  | Day 3 | 106.1 | 150.2 | 120.4 | 114.4 | 153.5 | 129.7 | 7.86% | 2.20% | 7.75% |
|  | Day 5 | 114.8 | 150.8 | 126.5 | 110.9 | 149.1 | 125.2 | 3.37% | 1.10% | 1.01% |
|  | Day 7 | 114.8 | 150.8 | 126.5 | 114.6 | 152.7 | 129.2 | 0.18% | 1.29% | 2.13% |
| M5 | Day 0 | 163.8 | 200.7 | 175.8 | 151.8 | 189.3 | 170.1 | 7.34% | 5.68% | 3.25% |
|  | Day 1 | 163.8 | 200.7 | 175.8 | 163.3 | 199.8 | 181.0 | 0.26% | 0.44% | 2.98% |
|  | Day 3 | 163.8 | 200.7 | 175.8 | 156.5 | 191.9 | 173.7 | 4.46% | 4.40% | 1.18% |
|  | Day 5 | 163.8 | 200.7 | 175.8 | 158.1 | 193.0 | 177.4 | 3.48% | 3.85% | 0.90% |
|  | Day 7 | 163.8 | 200.7 | 175.8 | 152.0 | 188.7 | 170.0 | 7.21% | 5.99% | 3.27% |

### *In vivo* flow data and data extraction

Pulsed wave (PW) Doppler velocity data was obtained at baseline, diagnosis day, and days 1, 3, 5, and 7 post diagnosis. Locations for PW Doppler data acquisition were positioned at the proximal aorta, the distal aorta, and at all arteries that are included in the 3D model: the celiac artery, left and right renal arteries, and superior mesenteric artery. PW Doppler data was analyzed with existing software (12, 13) at all outlets of the 3D model to extract the *in vivo* cycle averaged maximum velocity **v**_max_ at the centerline of the vessel.

Assuming a parabolic cross-sectional velocity profile and a circular cross-section, we determined the mean velocity at each outlet as 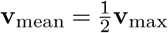. The cycle averaged flowrate *Q*_*i*_ at each outlet *i* was approximated as *Q*_*i*_ = **v**_mean,*i*_ ·*A*_*i*_, where *A*_*i*_ denotes the cross-sectional area of outlet *i*. Finally, we determined the flow ratios *q*_*i*_ as the ratio of total outflow (Table 2). In cases where the imaging data were not suitable for flowrate extraction, we prescribed data from a previous acquisition (indicated by an asterisk in Table 2).

**Table 2.** RCR–boundary condition parameters, identified by tuning to case specific data for the default anatomic models. *R*_*T*_ denotes the total resistance, *C*_*T*_ denotes the total compliance, and *k*_*d*_ is the ratio of distal to total resistance. The flow–splits *q*_*i*_ are presented as the ratio of total flow that exits through the respective vessel. The asterisk marks data that have not been available for the specific mouse/day combination and flow rates from the next available date were used to calculate the flow ratios.

|  | M2 |  |  |  |  |
| --- | --- | --- | --- | --- | --- |
|  | Day 0 | Day 1 | Day 3 | Day 5 | Day 7 |
| $R_T(\frac{g}{s\ mm^4})$ | 236.82 | 217.63 | 267.42 | 370.22 | 251.46 |
| $C_T(\frac{mm^4\ s^2}{g})$ | 1.26E-03 | 1.12E-03 | 5.61E-04 | 6.59E-04 | 7.35E-04 |
| $k_d(-)$ | 0.90 | 0.90 | 0.90 | 0.90 | 0.90 |
| $q_i(\text{celiac})$ | 0.09 | 0.10 | 0.07 | 0.13 | 0.09 |
| $q_i(\text{LRA})$ | 0.13 | 0.19 | 0.18 | 0.15 | 0.20 |
| $q_i(\text{RRA})$ | 0.03 * | 0.07 | 0.06 | 0.17 | 0.16 |
| $q_i(\text{SMA})$ | 0.16 | 0.24 | 0.16 | 0.24 | 0.27 |
| $q_i(\text{distal})$ | 0.59 * | 0.39 | 0.52 | 0.32 | 0.28 |
|  | M3 |  |  |  |  |
|  | Day 0 | Day 1 | Day 3 | Day 5 | Day 7 |
| $R_T(\frac{g}{s\ mm^4})$ | 135.17 | - | 203.52 | 287.73 | 322.55 |
| $C_T(\frac{mm^4\ s^2}{g})$ | 2.04E-03 | - | 9.31E-04 | 8.48E-04 | 6.03E-04 |
| $k_d(-)$ | 0.90 | - | 0.94 | 0.95 | 0.94 |
| $q_i(\text{celiac})$ | 0.19 | - | 0.19 | 0.18 | 0.13 |
| $q_i(\text{LRA})$ | 0.13 * | - | 0.25 | 0.16 | 0.07 |
| $q_i(\text{RRA})$ | 0.21 | - | 0.13 | 0.22 * | 0.08 |
| $q_i(\text{SMA})$ | 0.26 | - | 0.16 | 0.22 | 0.24 |
| $q_i(\text{distal})$ | 0.21 | - | 0.26 | 0.21 | 0.47 |
|  | M4 |  |  |  |  |
|  | Day 0 | Day 1 | Day 3 | Day 5 | Day 7 |
| $R_T(\frac{g}{s\ mm^4})$ | - | 485.26 | 291.50 | 296.80 | 307.86 |
| $C_T(\frac{mm^4\ s^2}{g})$ | - | 7.30E-04 | 8.33E-04 | 1.06E-03 | 9.70E-04 |
| $k_d(-)$ | - | 0.85 | 0.83 | 0.83 | 0.82 |
| $q_i(\text{celiac})$ | - | 0.21 | 0.17 | 0.10 | 0.06 |
| $q_i(\text{LRA})$ | - | 0.08 | 0.13 | 0.12 | 0.17 |
| $q_i(\text{RRA})$ | - | 0.14 | 0.06 | 0.14 | 0.10 |
| $q_i(\text{SMA})$ | - | 0.24 | 0.17 | 0.36 | 0.17 |
| $q_i(\text{distal})$ | - | 0.33* | 0.47 | 0.29 | 0.50 |
|  | M5 |  |  |  |  |
|  | Day 0 | Day 1 | Day 3 | Day 5 | Day 7 |
| $R_T(\frac{g}{s\ mm^4})$ | 337.35 | 417.48 | 296.13 | 262.41 | 411.36 |
| $C_T(\frac{mm^4\ s^2}{g})$ | 6.12E-04 | 6.39E-04 | 9.00E-04 | 8.09E-04 | 5.83E-04 |
| $k_d(-)$ | 0.88 | 0.93 | 0.94 | 0.96 | 0.93 |
| $q_i(\text{celiac})$ | 0.07 | 0.04 | 0.05 * | 0.04 | 0.08 |
| $q_i(\text{LRA})$ | 0.13 | 0.17 | 0.21 | 0.29 | 0.16 |
| $q_i(\text{RRA})$ | 0.25 | 0.28 | 0.06 | 0.13 | 0.27 |
| $q_i(\text{SMA})$ | 0.33 | 0.19 | 0.11 | 0.14 | 0.18 |
| $q_i(\text{distal})$ | 0.22 | 0.32 | 0.57 | 0.39 | 0.30 |

A time-resolved inlet velocity wave form was extracted at the centerline of the proximal aorta to prescribe the inflow boundary condition in the computational fluid dynamics (CFD) simulations.

### Computational model

#### Anatomic model construction and mesh generation

The anatomic models were segmented based on threedimensional B-mode US images. Color Doppler and EKV US modalities were additionally consulted in cases where intestinal gas reduced image quality by obscuring portions of the underlying abdominal aorta. The 3D model domain comprises the abdominal aorta and the following branching vessels: celiac artery, left and right renal arteries, and superior mesenteric artery (Figure 1a).

**Fig. 1.**
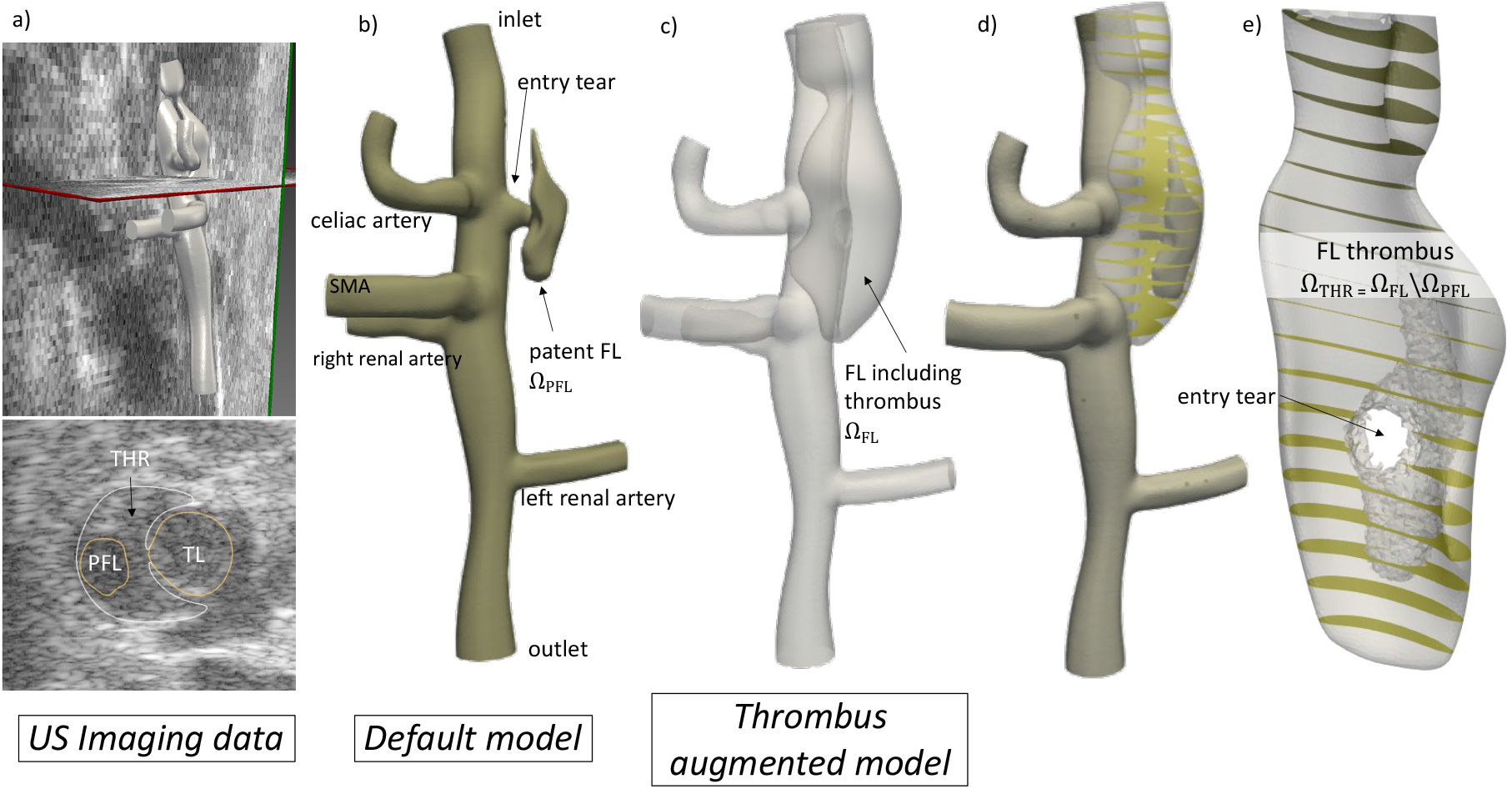
Specimen M3, Day7. a) Example of US imaging data: 3D view with *thrombus augmented model* (top) and axial plane (bottom) with segmentation of thrombus (THR), patent FL, and TL. b) *Default model* with patent FL Ω_PFL_, celiac artery (CA), superior mesenteric artery (SMA), right renal artery (RRA), and left renal artery (LRA). c) The *thrombus augmented model* combines patent and thrombosed FL to form domain Ω_FL_. d) Visual overlay of the default and thrombus augmented model. e) Thrombosed model domain Ω_THR_ = Ω_FL_ \Ω_PFL_ with entry tear.

Two model variants were created for each animal: In the *de-fault model*, regions of TL and FL, and the fenestrating tears between TL and FL were identified and incorporated in the anatomic models. Thrombosed regions were treated as impermeable to flow, and not included in the 3D model domain. In the *thrombus augmented model* (created for Day 7 post diagnosis only), thrombosed regions were treated as part of the fluid domain and included into the 3D model domain. No animal developed thrombus outside of the FL.

Image segmentation and model generation were performed in SimVascular (14), an open source cardiovascular flow modeling software, with supplemental editing performed in Meshmixer (Autodesk, Inc.). Three-dimensional unstructured tetrahedral meshes were subsequently created with the TetGen mesh generator (15), which is embedded within SimVascular.

#### Governing equations

Blood was modeled as an incompressible, Newtonian fluid and governed by the Navier–Stokes equations,

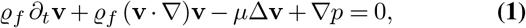

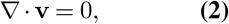

where **v** and *p* denote the velocity and pressure, respectively. We set *ϱ*_*f*_ = 1.06 g*/*cm^3^, *μ* = 0.04 g*/*cmsec. We treated the outer domain walls as rigid and assumed a no-slip boundary condition, **v** = 0 at the wall of the fluid domain. This assumption is supported by time-resolved imaging data which shows little movement of the vessel wall during the cardiac cycle.

#### Boundary conditions

We prescribed a pulsatile flow waveform with a parabolic velocity profile at the inlet of the computational domain informed by the time resolved flow rate data for each mouse and day (described above).

At the outlets we prescribed three-element Windkessel (RCR) boundary conditions, following the coupled multidomain method as described in (16, 17). Specimen and day specific flow ratios and pressure data were available and used as target values. Parameters for the RCR outlets were tuned to match the experimental observations within a 10% error margin.

The total resistance, compliance and the ratio between proximal and distal resistance for the respective simulations are given in Table 2. The total resistance *R*_*T*_ and capacitance *C*_*T*_ were distributed proportionally to the flow ratio at each outlet *i*: *R*_*T,i*_ = *R*_*T*_ */q*_*i*_ and *C*_*i*_ = *C*_*T*_ · *q*_*i*_. This distribution yields an approximation to the flow-splits as given in Table 2. The ratio of distal to proximal resistance is denoted by the constant value *k*_*d*_ ∈ (0, 1). Proximal and distal resistance at each outlet are then given by *R*_*d,i*_ = *k*_*d*_*R*_*T,i*_ and *R*_*p,i*_ = (1 − *k*_*d*_)*R*_*T,i*_. To reduce computational cost, we performed an initial RCR parameter tuning step on coarse meshes with a spatial resolution of 0.1 mm, resulting in meshes of less than 100,000 elements. Once the pressure targets for diastolic, systolic, mean and pulse pressure were closely approximated on the coarse meshes, we refined the tuning process on meshes with a spatial resolution of 0.06 mm and approximately 500,000 elements. This mesh size had been identified as sufficiently fine in a mesh independence analysis (see below). For the simulations on thrombus-augmented models, we applied RCR parameters from the corresponding default models. To achieve cycle-to-cycle periodicity, we observed an initialization period of at least 9 cardiac cycles, and reported simulation results from the 10^th^ or subsequent cardiac cycle, following (18).

#### Numerical discretization

The numerical simulations were performed with svFSI, a finite element solver included in SimVascular featuring linear elements for velocity and pressure and a generalized-*α* scheme for temporal discretization. For stabilization, the P1/P1 finite element space is augmented by weighted residuals, known as the Residual Based Variational Multiscale Method (RBVMS). The RBVMS has been validated in a wide range of cardiovascular simulations (17, 19). At the fluid outlets, backflow stabilization was applied (17).

### Mesh sensitivity analysis and temporal resolution

To identify a sufficiently small spatial resolution for the tetrahedral meshes, we performed a mesh independence study on model Mouse M4, Day5. We created 4 tetrahedral meshes, with varying meshsizes (Table 3). Pressure and wall shear stress (WSS) results were compared to simulation results on the finest mesh (h = 0.05 mm). We observed pressure and WSS errors below 3% on the two meshes with meshsize h = 0.1 mm and h = 0.75 mm. Errors further reduced to below 1% on the mesh with h = 0.6mm. Based on these results we set the mesh size for subsequent simulations to 0.6mm. Given the very small errors in WSS in the FL (all errors were below 10^−3^ Pa), we did not include FL WSS results in the mesh sensitivity analysis.

**Table 3.** Results of mesh refinement analysis in four meshes. We based the identification of a sufficiently small mesh size on the relative errors in pressure in the TL and FL (*e*^P,TL^ and *e*_P,FL_, respectively) and relative errors in TL WSS (*e*_WSS,TL_). All observed errors in FL WSS were below 1 mPa, and thus not evaluated for relative errors.

| mesh size (mm) | $h = 0.1$ | $h = 0.075$ | $h = 0.06$ |
| --- | --- | --- | --- |
| $e_{P,TL}$ | 1.5% | 0.9% | 0.2% |
| $e_{P,FL}$ | 1.4% | 0.8% | 0.2% |
| $e_{WSS,TL}$ | 2.8% | 2.3% | 0.9% |

The temporal resolution was chosen such that each cardiac cycle contained 1,000 time steps. For post-processing of the numerical results, detailed spatially resolved simulation results were saved every 20^*th*^ time-step.

### Thrombus volume

To quantify the evolution of thrombus volume and FL growth we measured the volume of patent FL, V_PFL_, and estimated thrombus volume V_THR_ by (i) creating a segmentation of the aorta including thrombus and patent FL, and then (ii) subtracting the segmented aorta from the default model. Additionally we determined the sum of both V_FL_ = V_PFL_ + V_THR_ volume over total FL volume,

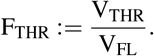

### Average velocity and pressure in the FL

We calculated the relative average velocity and pressure in the FL as

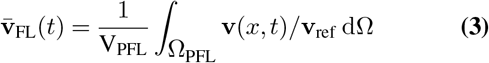

and

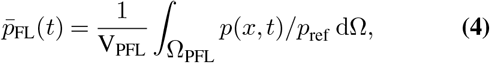

where **v**_ref_ and *p*_ref_ denote the reference velocity and pressure at the proximal, undissected aorta.

Cycle averaged quantities were computed as

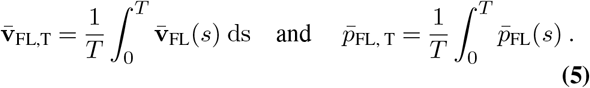

### Particle residence times

Regions of stagnant flow are prone to thrombus formation and growth. Particle residence times can be utilized to identify such regions and assist in understanding local variations in thrombus deposition, (20, 21). We quantified residence times (RT) following an approach suggested by (22) in which a non-discrete method to calculate RT is introduced in an Eulerian framework using the advection-diffusion equation. The resulting metric, *τ* (*x, t*), quantifies the time a fluid particle has been located inside a predefined region of interest, in our case the FL. RTs are calculated with an in-house finite element solver, described in (22). The calculation of residence times was performed with converged, periodic flow results. In the literature, typical runtimes for the particle residence time solver were reported as 6 cardiac cycles or a maximum of 12 sec, respectively (22, 23). Since runtime was not an issue in our application, we computed residence times over a duration of 20 cardiac cycles.

### Endothelial cell activation potential (ECAP), platelet activation potential (PLAP), and thrombus formation potential (TFP)

As a measure for the degree of ‘thrombogenic susceptibility’ of the vessel wall, (24) defined the endothelial cell activation potential (ECAP),

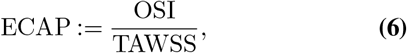

which takes into account that wall regions of high oscillatory shear index (OSI) and low TAWSS are receptive to the deposition of thrombus.

Platelet activation potential (PLAP) is a hemodynamic marker that quantifies cumulative shear along pathlines and thus serves as a index for the mechanical activation of platelets in the fluid field (24, 25). It is defined as

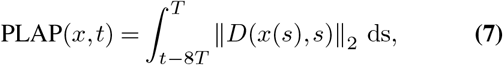

where ‖ · ‖_2_ denotes the Frobenius norm and 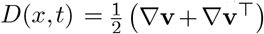 denotes the symmetric part of the velocity gradient.

To calculate PLAP we follow the suggested approach by (24) and build upon previous work by (26): A fourth order RungeKutta scheme is employed to determine trajectories of massless particles along the pre-computed transient flow field. Similar to (24), we tracked particles backwards in time and used seed locations close to the surface of the vessel wall. This approach enabled us to focus on particles that eventually come close to the endothelial layer and FL surface. Target particles were located at cell centers within a prescribed threshold of h = 0.1 mm from the vessel surface. We considered five injection times per cardiac cycle and followed the particles until either (i) the maximum integration time of 8 cardiac cycles has elapsed, (ii) the particle has left the 3D computational domain, or (iii) the particle is stagnating for an entire cardiac cycle. Results were subsequently averaged across the different injection times.

TFP combines ECAP and PLAP,

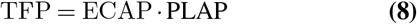

and may be used to identify surface regions where a susceptible endothelium may come into contact with activated platelets (24).

## Results

### Anatomic changes over time

Three specimens (M2, M3, and M4 the naming is adopted from (10)) presented with a steady morphology of TL and FL during the time frame of observation. M2 and M3 both developed a single FL with a single visible entry tear and M4 developed a single FL with two distinct visible fenestrations. In an additional animal (M5), the morphology of the dissection changed during the time frame of observation (Figure 2). In detail, on the day of diagnosis (day 0) we detected two separate false lumina: the proximal FL was connected to the TL via a single tear and the distal FL had a distinct entry and exit tear. On day 1, we observed two false lumina, which were connected to the TL via a single tear. The topology day 3 resembled day 0. On day 5 post diagnosis, we observed 3 false lumina, two of which had a single tear. The most distal FL had two tears for a total of 4 openings. On day 7 we observed a single FL with three distinct tears. This rapidly evolving morphology suggests a highly complex remodeling process for M5.

**Fig. 2.**
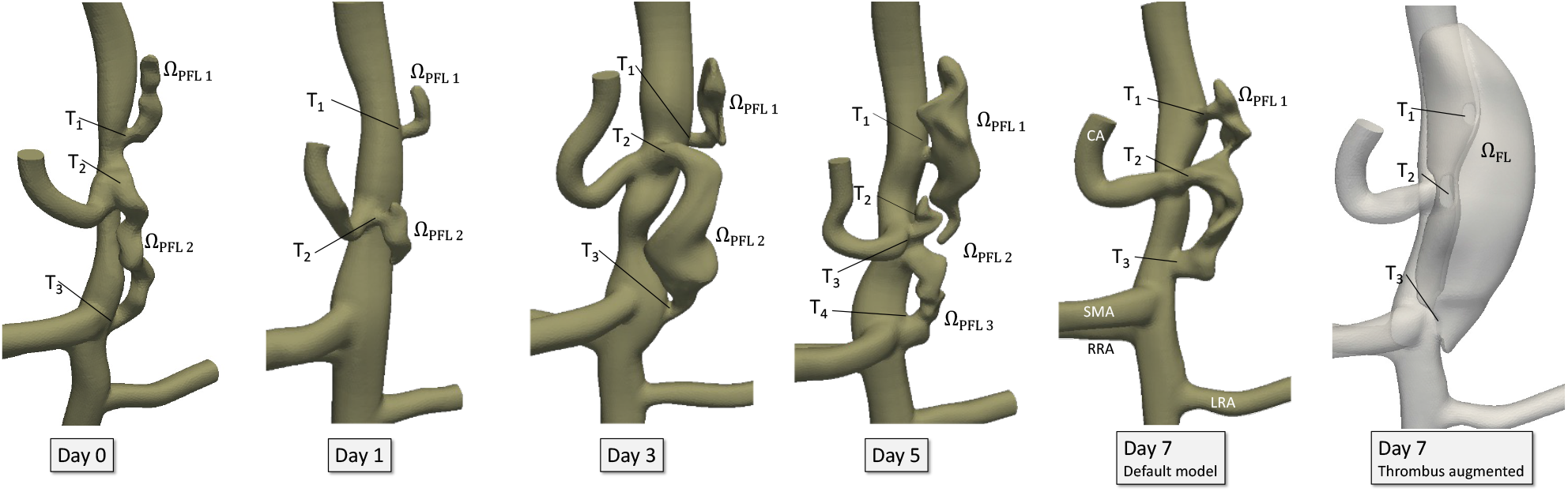
3D model of Mouse M5 during the time frame of observation.

### Thrombus volume over time

All specimens presented with a substantial amount of thrombus on the day of diagnosis. The total FL volume V_FL_ remained relatively stable at approximately 10 mm^3^ for mouse models M2, M3 and M5, and reached up to 20 mm^3^ in specimen M4. Model M4 also displayed the largest patent FL volume V_PFL_ and the largest V_FL_ and V_PFL_ variation over time (Fig. 3).

**Fig. 3.**
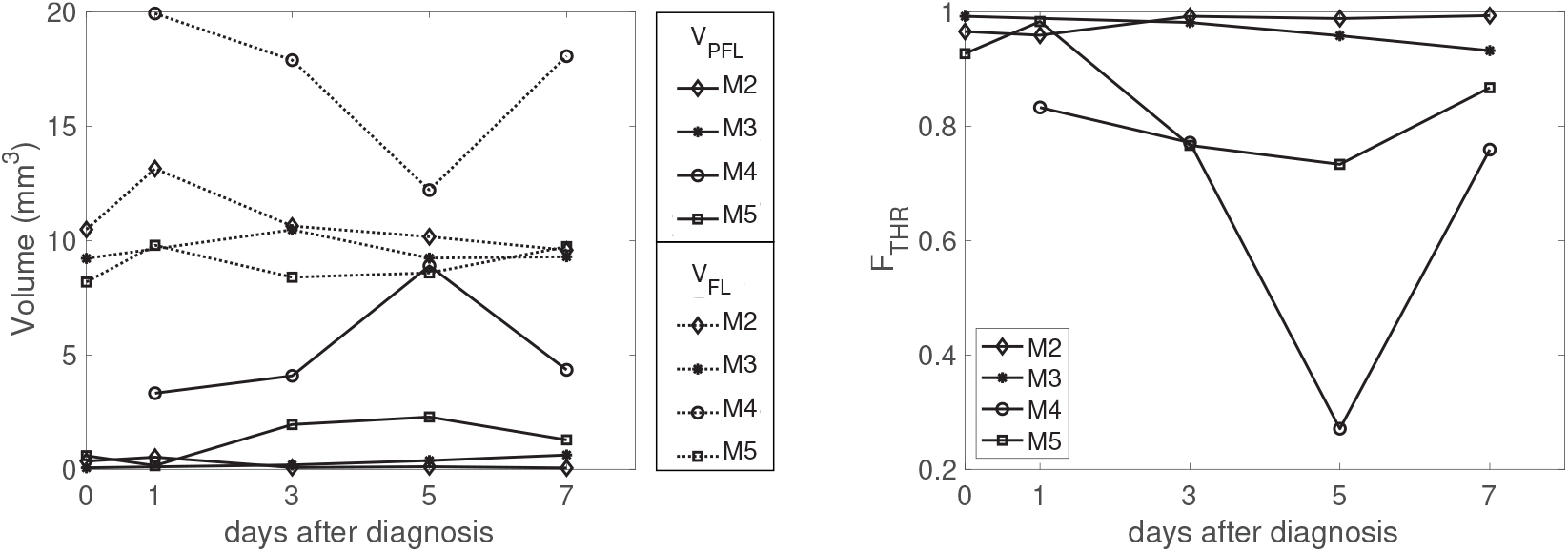
Volume of patent FL V_PFL_ and volume of FL including thrombus V_FL_, and thrombus ratio F_THR_ remain relatively stable in specimen M2, M3, M5. Thrombus ratio F_THR_ remains above 95% in M2 and M3, and varies substantially in M4.

The ratio of thrombus volume over total FL volume, F_THR_, remained large in M2 and M3 (F_THR_ *>* 0.93) on all observation days and F_THR_ averages above 0.96 for both mice. Both cases had false lumina that are connected to the TL with a single tear.

Models M4 and M5 displayed a greater variability of F_THR_, with F_THR_ ∈ [0.27, 0.83] for M4 and F_THR_ ∈ [0.73, 0.98] for M5 and with averages of 0.65 and 0.86 respectively. These specimens had two or more fenestrations in the dissected region of the aorta.

### Velocity and pressure in FL

The smallest relative mean velocities 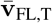 were observed in false lumina with a single fenestration and the 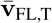 remained below 0.016, 0.005, and 0.035 in M2, M3 and M5, respectively (Table 4, Figure 4). In false lumina with two or more tears, 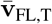 displayed slightly larger values with a relative mean velocity 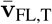 below 0.064 and 0.099 in M4 and M5, respectively (Table 4, Figure 4). Combining data from all false lumina with a single tear, 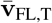 had a mean value of 0.006 with standard deviation 0.009. For lumina with two tears, 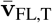 had a mean of 0.045 with standard deviation 0.035, Figure 4. Simulation results did not reveal noticeable pressure differences between TL and FL and values for 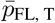 deviated less than 2.7% from the mean pressure at the inlet of the computational domain (Table 4).

**Table 4.** Simulation results of relative mean velocity (i.e. the velocity was averaged over the false lumen and cardiac cycle and divided by the reference velocity at the inlet). The days are counted post diagnosis. The FL are numbered from proximal to distal.

| M2 |  |  |  |  |  |  |
| --- | --- | --- | --- | --- | --- | --- |
|  | Day 0 | Day 1 | Day 3 | Day 5 | Day 7 | Day 7 THR |
| $\bar{v}_{FL,T}$ | 1.408E-03 | 2.216E-04 | 4.240E-05 | 8.847E-05 | 1.620E-02 | 3.460E-04 |
| $\bar{p}_{FL,T}$ | 0.981 | 0.988 | 0.992 | 0.995 | 0.992 | 0.992 |
| Number of tears | 1 | 1 | 1 | 1 | 1 | 1 |

| M3 |  |  |  |  |  |  |
| --- | --- | --- | --- | --- | --- | --- |
|  | Day 0 | Day 1 | Day 3 | Day 5 | Day 7 | Day 7 THR |
| $\bar{v}_{FL,T}$ | 2.698E-03 | - | 4.846E-03 | 3.070E-03 | 3.932E-03 | 3.658E-04 |
| $\bar{p}_{FL,T}$ | 0.993 | - | 0.995 | 0.995 | 0.996 | 0.994 |
| Number of tears | 1 | - | 1 | 1 | 1 | 1 |

| M4 |  |  |  |  |  |  |
| --- | --- | --- | --- | --- | --- | --- |
|  | Day 0 | Day 1 | Day 3 | Day 5 | Day 7 | Day 7 THR |
| $\bar{v}_{FL,T}$ | - | 4.305E-02 | 2.777E-02 | 6.408E-02 | 5.668E-03 | 2.296E-03 |
| $\bar{p}_{FL,T}$ | - | 0.987 | 0.982 | 0.978 | 0.992 | 0.992 |
| Number of tears | - | 2 | 2 | 2 | 2 | 2 |

| M5 |  |  |  |  |  |  |
| --- | --- | --- | --- | --- | --- | --- |
|  | Day 0 | Day 1 | Day 3 | Day 5 | Day 7 | Day 7 THR |
| $\bar{v}_{FL,TFL1}$ | 8.211E-03 | 1.001E-02 | 1.207E-04 | 7.870E-04 | 4.082E-02 | 4.919E-02 |
| $\bar{v}_{FL,TFL2}$ | 9.938E-02 | 1.965E-03 | 6.774E-02 | 3.480E-02 | - | - |
| $\bar{v}_{FL,TFL3}$ | - | - | 5.094E-03 | - | - | - |
| $\bar{p}_{FL,TFL1}$ | 0.983 | 0.994 | 0.995 | 0.995 | 0.994 | 0.995 |
| $\bar{p}_{FL,TFL2}$ | 0.973 | 0.993 | 0.993 | 0.992 | - | - |
| $\bar{p}_{FL,TFL3}$ | - | - | - | 0.993 | - | - |
| Number of tears FL1 | 1 | 1 | 1 | 1 | 3 | 3 |
| Number of tears FL2 | 2 | 1 | 2 | 1 | - | - |
| Number of tears FL3 | - | - | - | 2 | - | - |

**Table 5.** Simulation results of relative mean velocity (divided by reference velocity at inlet, averaged over FL volume, averaged over cardiac cycle) and relative mean pressure.

| Mouse M2 ( 1 tear) |  |  |  |
| --- | --- | --- | --- |
|  | Day 7<br>default model | Day 7 THR<br>augmented model | Day 7<br>Thrombus minus patent |
| $\bar{v}_{FL,T}$ | 1.620E-02 | 3.460E-04 | 1.033E-04 |
| $\bar{p}_{FL,T}$ | 0.992 | 0.992 | 0.992 |

| Mouse M3 (1 tear) |  |  |  |
| --- | --- | --- | --- |
|  | Day 7<br>default model | Day 7 THR<br>augmented model | Day 7<br>Thrombus minus patent |
| $\bar{v}_{FL,T}$ | 3.93E-03 | 3.66E-04 | 1.24E-04 |
| $\bar{p}_{FL,T}$ | 0.996 | 0.994 | 0.996 |

| Mouse M4 (2 tears) |  |  |  |
| --- | --- | --- | --- |
|  | Day 7<br>default model | Day 7 THR<br>augmented model | Day 7<br>Thrombus minus patent |
| $\bar{v}_{FL,T}$ | 5.67E-03 | 2.30E-03 | 8.58E-04 |
| $\bar{p}_{FL,T}$ | 0.992 | 0.992 | 0.992 |

| Mouse M5 (3 tears) |  |  |  |
| --- | --- | --- | --- |
|  | Day 7<br>default model | Day 7 THR<br>augmented model | Day 7<br>Thrombus minus patent |
| $\bar{v}_{FL,T}$ | 4.082E-02 | 4.919E-02 | 4.307E-02 |
| $\bar{p}_{FL,T}$ | 0.994 | 0.995 | 0.995 |

**Fig. 4.**
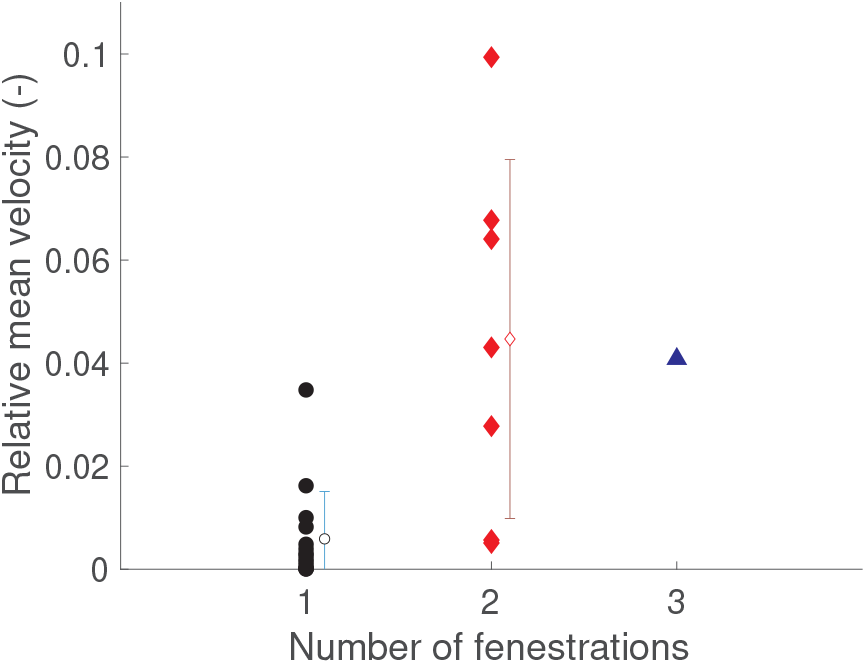
Relative averaged velocity 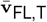 with mean and SD in FL grouped with respect to the number of fenestrations.

### Particle residence times

FL with a single tear (as seen in models M2 and M3, proximal FL in M5) exhibited elevated levels of *τ* throughout the FL, Fig 5. The calculated residence times reach values of approximately 2 sec, which corresponds approximately to the total runtime of 20 cardiac cycles, indicating stagnant flow in the FL. Similarly, in the end-regions of FL with multiple tears (M4, distal FL in M5), residence times reach values of *τ* ≈ 2 sec indicating near total stagnation in those regions.

**Fig. 5.**
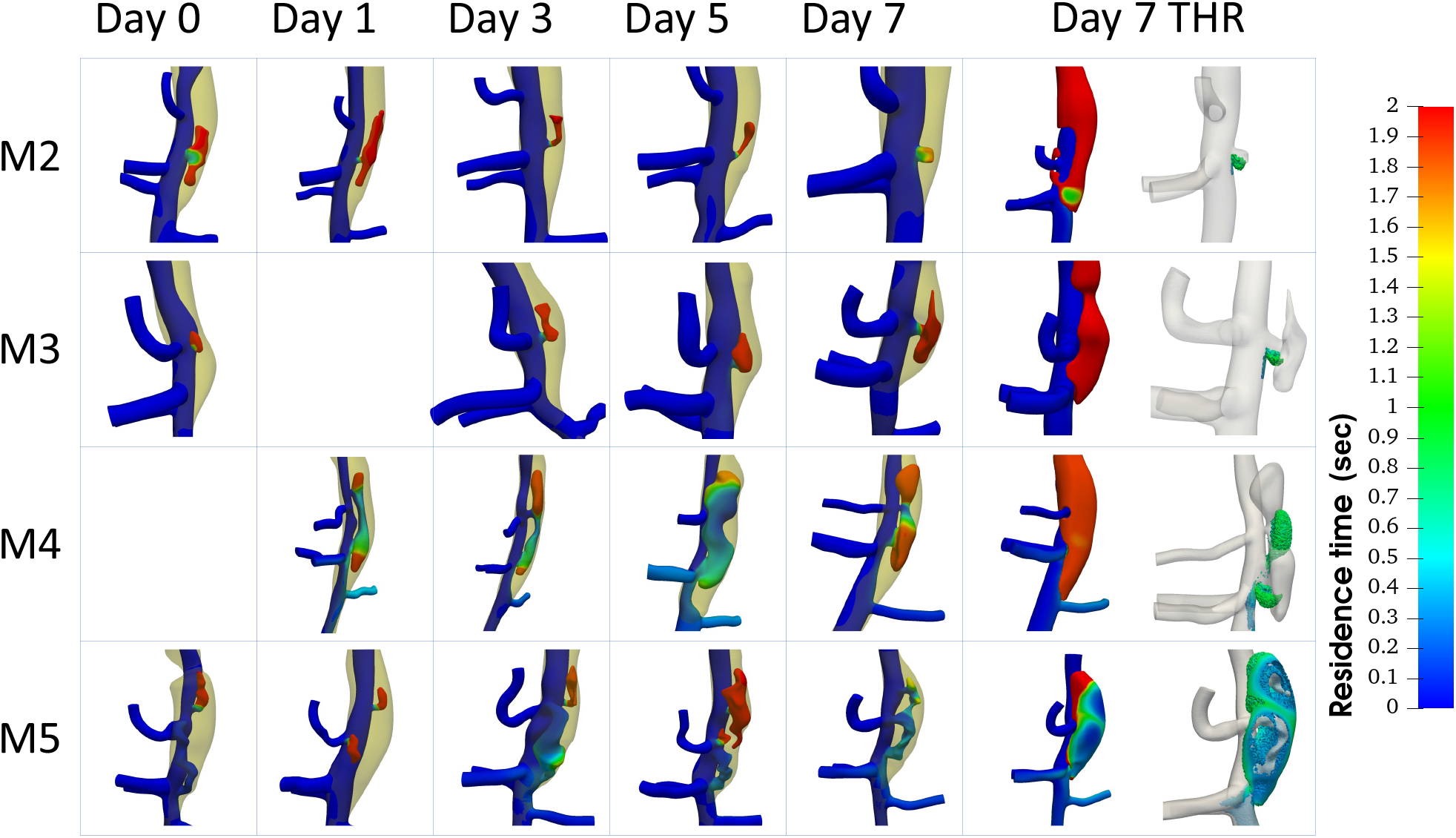
Particle Residence Time *τ* (in sec) after 20 cardiac cycles. A residence time of 2 seconds indicates stagnant flow. We show an overlay of thrombosed regions for days all five scan days. On day 7 residence times are calculated in the thrombus augmented models (‘Day 7 THR’). The right panel shows an overlay of residence times *τ* ∈ [0.3, 0.9] with patent FL.

### Endothelial cell activation potential (ECAP)

Without exception, we found that ECAP values were several orders of magnitude lower on the TL outer wall and branch vessel surfaces compared to FL with a single tear. Typically, ECAP remained below 0.5 on the TL outer wall and branch vessel surfaces. Especially in FL with a single tear, ECAP values were substantially increased, and typically exceeded ECAP = 1,000 (Figure 6). In contrast, ECAP values were low on the FL surface of M5, Day 7, a FL with three connected tears allowing for increased flow through the FL. Similarly, in FL with two tears, ECAP values typically remained below 0.5 in the regions connecting the tears, see e.g. M4 in Figure 6. ECAP ranges varied substantially between specimen and within each specimen over time. Maximum ECAP in M2 reached 9.2E6 on Day 3, and max ECAP = 12 on Day 7, on which the FL was reduced to a bulge. In M3, maximum ECAP reached 3.6E7 in the thrombus augmented model on Day 7 and max ECAP = 5.9E3 on Day 0 which again corresponded to the FL with the smallest extend. In M4, maximum ECAP reached 1.1E6 in the thrombus augmented model on Day 7 and max ECAP = 58 on Day 5. In M5, maximum ECAP reached 2.5E7 on Day 3, and max ECAP = 401 on Day 7 where the FL was connected by 3 tears to the TL.

**Fig. 6.**
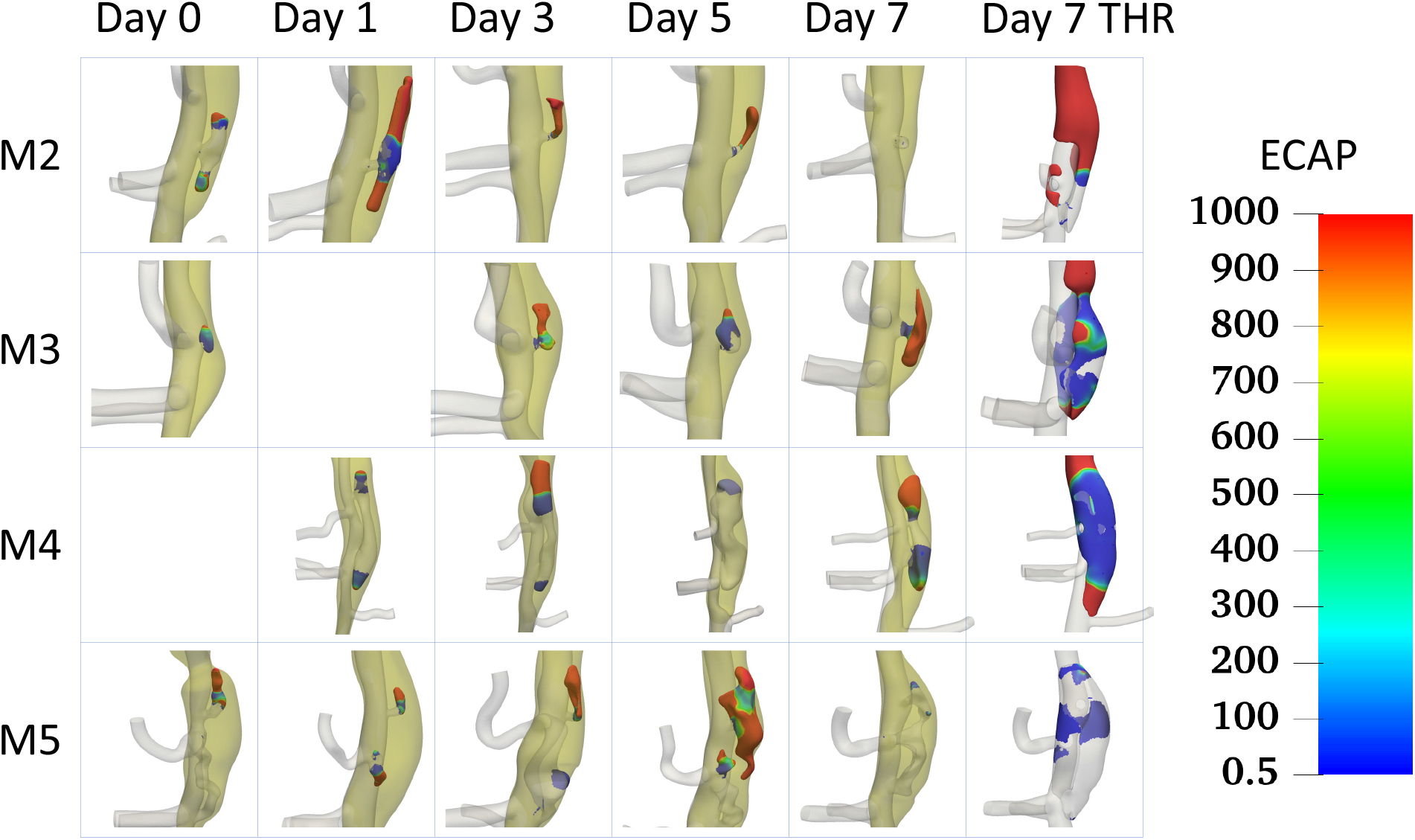
Regions of ECAP with ECAP *>* 0.5. For simulations on the default models, the regions of thrombus are shown. ECAP remains below 0.5 on the outer wall of TL and undissected branch vessels. Especially in FL with a single tear, (M2, M3 and some FL in M5) ECAP values are increased.

### Platelet activation potential

The lowest PLAP values were observed in FL with a single tear or regions of stagnant flow and we typically observed PLAP *<* 0.1 in those regions. Outside of those regions of stagnant flow, PLAP was found to be substantially larger with values exceeding PLAP *>* 3000. The results are displayed in Fig. 7 for Model M5, a model that consists of FL with single and multiple tears.

**Fig. 7.**
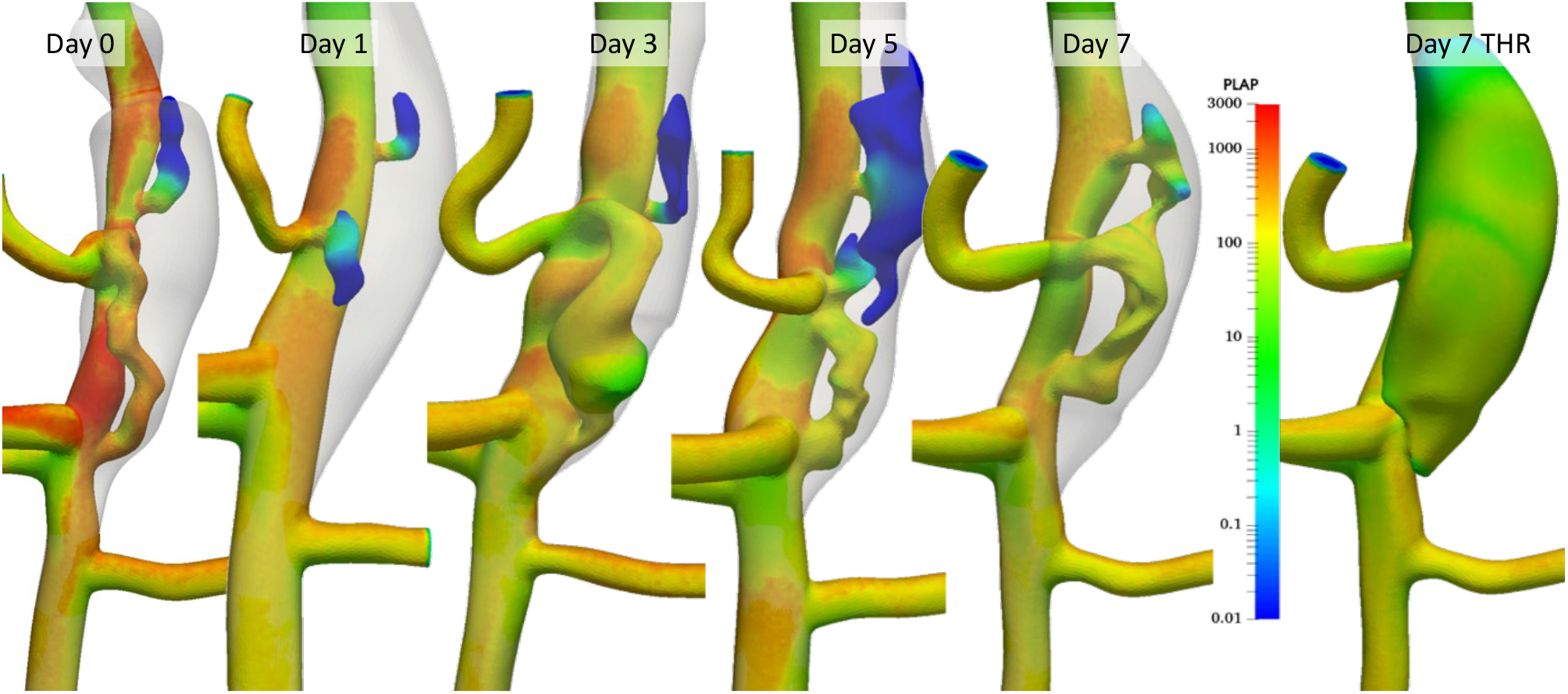
PLAP in model M5. Lowest values of PLAP are present in FL with a single connecting tear to the TL. Note the logarithmic lookup-table.

### Thrombus formation potential (TFP)

TFP is highest on the FL surface, especially in regions of stagnant flow. The range of TFP results varies substantially and we used a logarithmic lookup table to visualize the results in Figure 8. We also find regions of elevated TFP on the surface of the TL, for example in proximity to branch vessel ostia.

**Fig. 8.**
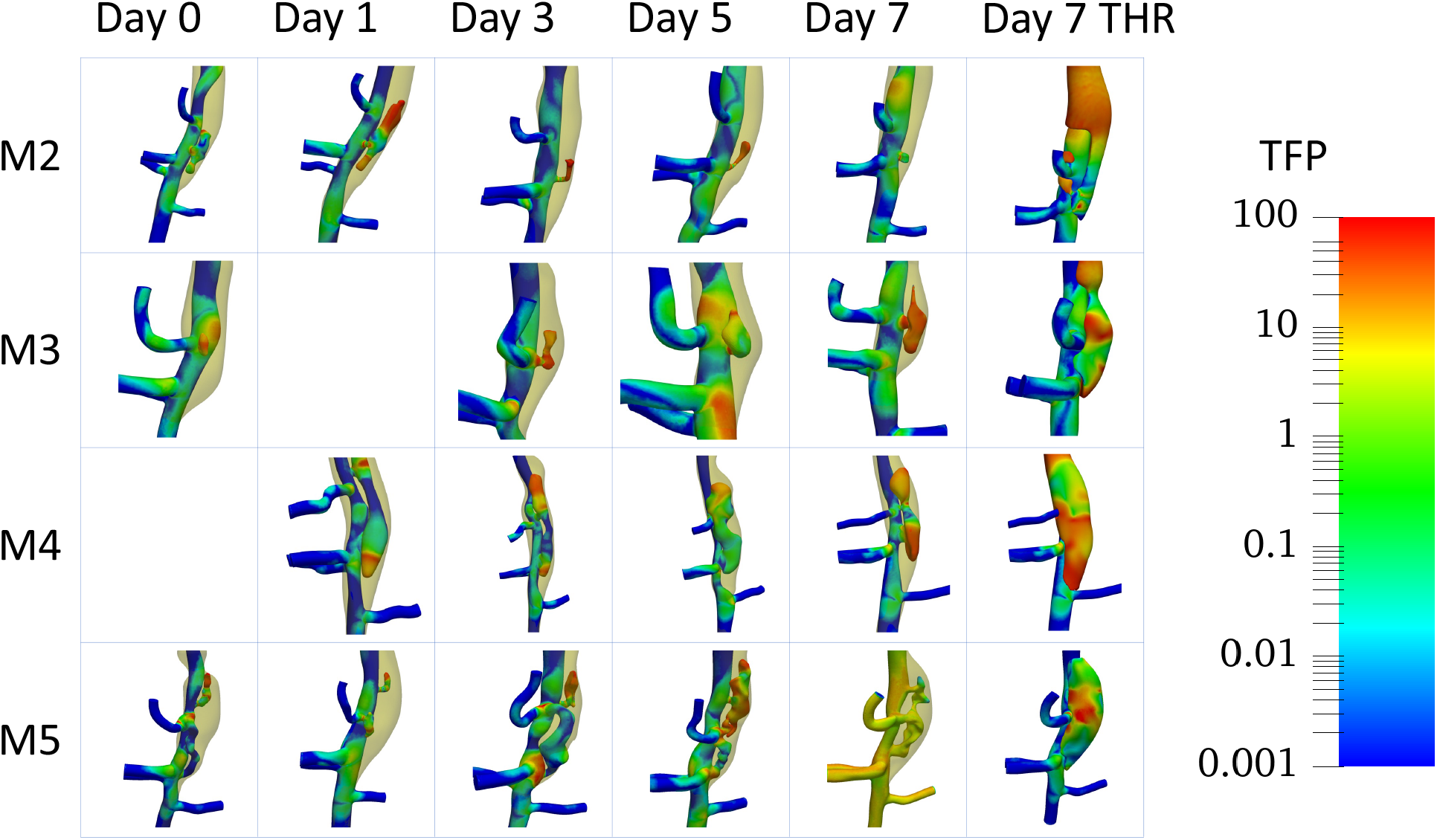
Thrombus formation potential (TFP). Note the logarithmic color scalebar.

### Comparison of mean velocity and pressure in default and thrombus augmented models

In addition to the *default models* discussed above, we also generated *thrombus augmented models* for day 7 in each mouse, in which the thrombosed regions are removed to mimic the pre-thrombus condition. Running the fluid dynamic simulations on these augmented geometries allows us to compare hemodynamic parameters in regions of the FL where thrombus is present compared to those regions where the FL remains patent. To this end we denote the domain of the patent FL in the default model by Ω_PFL_, the domain of the FL (patent and thrombosed) in the augmented model by Ω_FL_ and the difference of the two, i.e. the thrombosed regions as Ω_THR_ := Ω_FL_ \ Ω_PFL_, see Figure 1.

In all animals except for M5, the mean reference velocity in Ω_THR_ is smaller than the reference velocity in Ω_PFL_. In specimens M4 and M5, 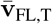 is of the same order of magnitude in Ω_PFL_ and Ω_FL_. Both specimen have two or more tears supplying the FL with fluid, as well as the lowest ratio of THR over FL (Figure 3). In contrast, in models M2 and M3, the mean velocity 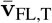 is of the same order of magnitude in Ω_FL_ and Ω_THR_. Both of these specimen have a single tear connecting the FL to the TL as well as the highest ratios of F_THR_ (Figures 3, 9). Comparing results for averaged pressures in the FL of all specimens, we find that 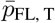 is similar in Ω_FL_ and Ω_THR_ with differences of less than 0.2% (Table 4).

**Fig. 9.**
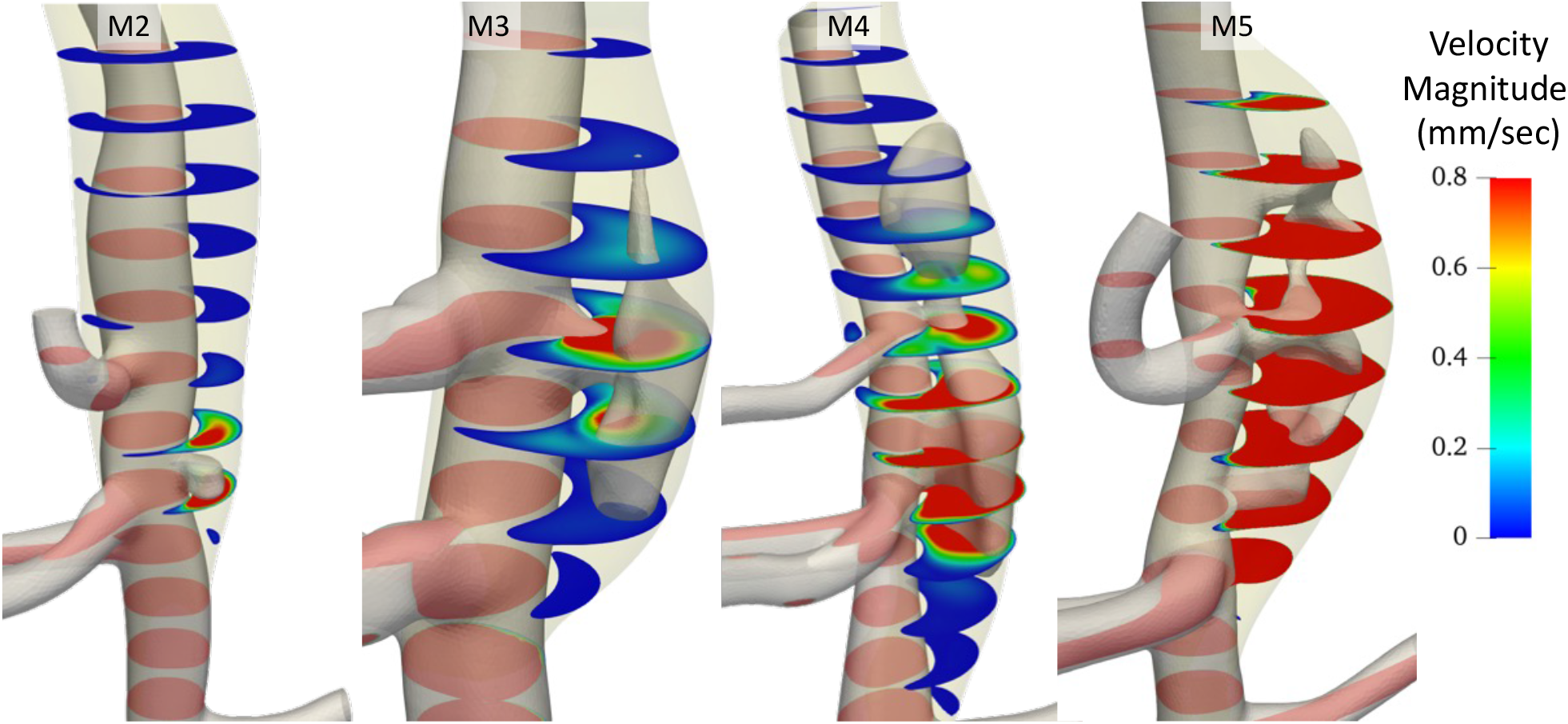
The colored cross-sections show velocity magnitude in the augmented models (day 7). The patent FL region is shown as an overlay. Models M2 abd M3 have a single connection from FL to TL, models M4 and M5 have multiple tears.

## Discussion

A particularly surprising observation of this murine study was the rapid evolution of thrombus volume and FL morphology over time. From clinical observations in humans we expect a gradual variation of thrombus volume in the FL. In these mice, thrombus varied substantially over the course of 7 days after onset of aortic dissection until the mice were euthanized. One mouse (M5) displayed a particularly interesting evolution in which even the number of FL and connecting openings varied on a daily basis. These observations could be attributed to the highly dynamic nature of these lesions or the challenges associated with accurately identifying thrombosed versus patent FL with *in vivo* US. In humans, (2) studied aortic dissections in a longitudinal study (median follow-up time of 4 years) and reported an increase of secondary tears or communications between true and false lumen over time. The authors explained this with an improved detectability of preexisting small tears over time within an increasingly thicker and less mobile dissection membrane. The proximal and distal extent of the false lumen and the location of the primary intimal tear were stable over time, suggesting dynamic differences between experimental and clinical aortic dissections.

Another interesting distinction between murine and human aortic dissection is the immediate presence of false lumen thrombosis in all four mice on the day of diagnosis. Per study design, the time-window between onset of aortic dissection and first US imaging did not exceed 48 hours, but naturally the exact time of onset is unknown. All mice presented with a thrombus ratio in the false lumen of more than 80% on their day of diagnosis. Contrastingly in humans, (1) and (3) report presence of thrombus in 43.3% and 56% of their patient cohorts, respectively. Thus the difference in intramural thrombus prevalence may be due to hemodynamic differences between mice and humans or potentially mouse model specific, and should be considered when translating finding from experimental aortic dissection studies.

In this study, observations in the four mice support the notion of a link between the extent of thrombus and the number of connecting tears in the respective false lumen. In FL with a single fenestration, more than 93% of FL volume was thrombosed. In FL with two or more entry tears 27% 93% of FL volume was thrombosed, with large day-to-day variations of up to 50% of the FL volume. This observation corresponds to our findings regarding the presence and extent of stagnant flow and corresponding long residence times in FL with a single tear. In FL with multiple tears, residence times were typically increased in regions proximal to the first tear and distal to the distal tear. Altogether, these results indicate that flow stagnation is the main driver of thrombosis in experimental murine AAAs. As a consequence of stagnant flow, TAWSS values are decreased substantially, leading to large ECAP values in the respective regions. Our findings suggest that ECAP appears to be mainly driven by low TAWSS values.

Another relevant metric in thrombosis, PLAP, provides a measure for the cummulative shear stress that a particle has experienced, which in turn provides a metric for the activation of platelets. If such a shear activated platelet reaches a surface with increased ECAP, one assumes an increased receptivity to thrombus deposition or TFP. In the investigated models, however, we found only small values of PLAP on the FL surface and we observed shear activated platelets predominantly on the TL surface. These findings align with the presence of reduced velocity gradients in regions of slow or stagnant flow, which are often close to the FL surface. Despite the low PLAP values in proximity to thrombus, the derived metric TFP in turn is elevated on the FL surface, a result that is likely caused by the substantially increased ECAP values in those regions. However, we also found elevated TFP values on TL surface, where none of the investigated mice developed thrombus. This suggests that shear activation of platelets plays a subordinate role in the present thrombus deposition and growth, and that flow stagnation is instead the main driver of thrombus deposition.

In summary, the evolution of thrombus volume in the investigated mice remained at constant high ratios in the FL with a single opening, and varied in cases with more than one FL fenestration. While residence times were consistently long in FL with largest thrombus ratio, thrombus deposition or growth did not correlate with hemodynamically derived metrics such as ECAP, PLAP, or TFP in this experimental dis-section model. Additional work is needed to identify other metrics that can be used to predict dissection expansion and intramural thrombus evolution.

## Limitations

One limitation of the current study is the number of investigated specimen. We considered four mice that varied widely in severity and represent mild, moderate, and severe dissections. Challenges in generating geometric models occurred due to our reliance on *in vivo* US imaging data, which possesses lower image to noise ratio than other *ex vivo* imaging approaches. A translation of our findings to human aortic dissections, amongst others, might be limited by potential differences in the these experimental dissections. Fore example, human aortic dissections typically have two or more connecting tears between the false and true lumen, and thrombus is present in a subset, whereas all investigated murine models had a partially thrombosed false lumen and several animals had a false lumen with only a single fenestration.

## ACKNOWLEDGEMENTS

This work used the Extreme Science and Engineering Discovery Environment (XSEDE), (27), on cluster Stampede2 at UT Austin, through allocation of P.I. Alison Marsden. XSEDE is supported by National Science Foundation grant number ACI-1548562. This work used the Stanford Research Computing Center (SRCC). Funding was also provided by the American Heart Association to Craig Goergen (14SDG18220010).

## Notes

### Competing Interest Statement

The authors have declared no competing interest.

